# SERBP1 is a master regulator of ribosome interactions

**DOI:** 10.64898/2026.08.26.747346

**Authors:** Nicolle A. Rosa-Mercado, Elif S. Hearn, Lukasz Szyrwiel, Kate L. Schole, Juri Rappsilber, Rachel Green

## Abstract

Ribosome function depends on interactions with diverse proteins whose activities determine translational efficiency, impose quality control and activate signaling pathways. These many competing activities must in turn be regulated. SERBP1 is an abundant cellular factor that interacts with ribosomes at multiple functionally critical sites on both dormant and active ribosomes. Here, we perform mass spectrometry across sucrose gradients in untreated and stressed cells and find that SERBP1 modulates ribosome interactions with many factors involved in processes including mRNA degradation, translational control, ribosome quality control and ribosome degradation. We then define the role of SERBP1 in protection of ribosomes against selective 40S degradation in the context of mTOR inhibition through its competition with the E3 ligase RNF10 and the atypical kinase RIOK3. Our work reveals SERBP1 as a key player in maintaining ribosome subunit balance and more broadly in the regulation of diverse ribosome activities.

## Introduction

While ribosomes are best known for their role in translating mRNAs into the encoded polypeptides, recent work has begun to reveal a broader role for ribosomes in reporting on cellular dysfunction and in functioning as a platform for the initiation of ribosome-mediated quality control and signaling pathways.^1^ Many key factors have been identified that play roles in these distinct outcomes, but an understanding of their interplay and their regulation is incompletely understood. Moreover, the molecular players involved in determining the balance between translation elongation, ribosome-mediated quality control, and the ribosome-mediated activation of cellular stress pathways like the ribotoxic stress response (RSR) and the integrated stress response (ISR) remain unknown.

Key regulatory factors are likely to be abundant cellular proteins that interact with ribosomes. Inactive ribosomes can be bound by hibernation factors including SERBP1, CCDC124, and IFRD2,^2–4^ but the broader role that these factors play on ribosomes remains poorly understood in human cells. SERBP1 (and its yeast homologue Stm1) has long been studied in the context of dormant ribosomes, where it binds to the vacant mRNA channel on non-translating 40S subunits.^2,3^ SERBP1 has also been shown to prevent ribosome degradation during mTOR inhibition, reportedly through mechanisms dependent on hypophosphorylation of SERBP1.^5^ However, the molecular mechanisms underlying SERBP1-mediated ribosome protection remain unknown. Because SERBP1 is highly abundant and approximately stoichiometric to ribosomes,^6^ it is unlikely that all copies of SERBP1 are bound to dormant ribosomes at a given time, especially in a cell that is robustly translating its mRNAs. Consistent with this idea, increasing evidence suggests that SERBP1 can bind to other positions on ribosomes compatible with active translation,^7–9^ raising the possibility that SERBP1 plays a more complex role beyond simply stably binding to dormant ribosomes. These recent studies show that SERBP1 has multiple “handles” on the ribosome: its N-terminus binds near the H79 helix of the 28S rRNA on the 60S, while different regions of the C-terminus interact with the 40S in the mRNA channel and in a hydrophobic pocket on the ribosomal protein RACK1.^9,10^ Recent work has shown that this hydrophobic pocket on RACK1 is also bound by the ribosome collision-responsive ZAKα kinase,^10^ which serves as the sensor for the ribotoxic stress response,^11–13^ as well as by LARP4.^14^ Moreover, the interaction between RACK1 and ZAK is antagonized by SERBP1 binding, hindering ZAK activation.^10^ These observations suggest that there may exist an even broader interactome at this site.

SERBP1 has been previously implicated in preventing ribosome degradation.^5^ Ribosome levels are maintained by tight regulation of ribosome biogenesis and degradation. The mammalian target of rapamycin (mTOR) impacts both processes, acting as a sensor for growth signals and nutrient levels.^15^ Cellular stress conditions leading to mTOR inhibition negatively impact cellular growth by decreasing translation, decreasing ribosome biogenesis and increasing ribosome degradation. In particular, many stress conditions that lead to ribosome stalling, as well as mutations in the rRNA that affect the decoding center or even 60S biogenesis defects,^16^ can lead to selective 40S degradation.^17–19^ 40S degradation in these settings relies on activation of the E3 ligase RNF10, which ubiquitylates the small subunit proteins uS3 and uS5. These ubiquitylations can be removed by the deubiquitylase USP10,^20,21^ but persistent ubiquitylation is recognized by the atypical kinase RIOK3.^17–19^ Binding of RIOK3 leads to the recruitment of the terminal uridylyl transferase TUT7 to uridylate the 18S rRNA, resulting in an oligo(U) tail that is then recognized by the exonuclease DIS3L2, which degrades the 18S rRNA.^22^ Despite our increasing understanding of the mechanisms that lead to stress-induced 40S degradation, our knowledge regarding how ribosomes are protected from sporadic degradation by these pathways remains incomplete.

Here we begin with a proteomic approach to define the breadth of the ribosome interactome that is regulated by SERBP1, identifying diverse factors involved in ribosome quality control (RQC), translational regulation, mRNA degradation, and selective 40S degradation. Focusing on a specific example, we then establish how SERBP1 binding to ribosomes protects 40S subunits from ubiquitylation and subsequent degradation through the RNF10/RIOK3 pathway. Together our data reveal SERBP1 as a key regulator of subunit balance and more generally of diverse ribosome interactions.

## Results

### SERBP1 binds translating ribosomes but does not impact elongation rates

We first used sucrose gradient sedimentation and western blotting for SERBP1 to determine whether SERBP1 is bound to polysomes in normally growing HEK-293T cells and in stressed cells treated with the mTOR inhibitor torin-1 (TOR) (Figure 1A). Despite the broad impact on translation and the overall increase in monosomes caused by mTOR inhibition, SERBP1 binds ribosomes robustly and its distribution across the gradient simply mirrors that of all ribosomes as followed by ribosomal protein eS24 (Figure 1A). Together with previous literature, these data suggest that SERBP1 can bind ribosomes regardless of their translational state.

**Figure 1:**
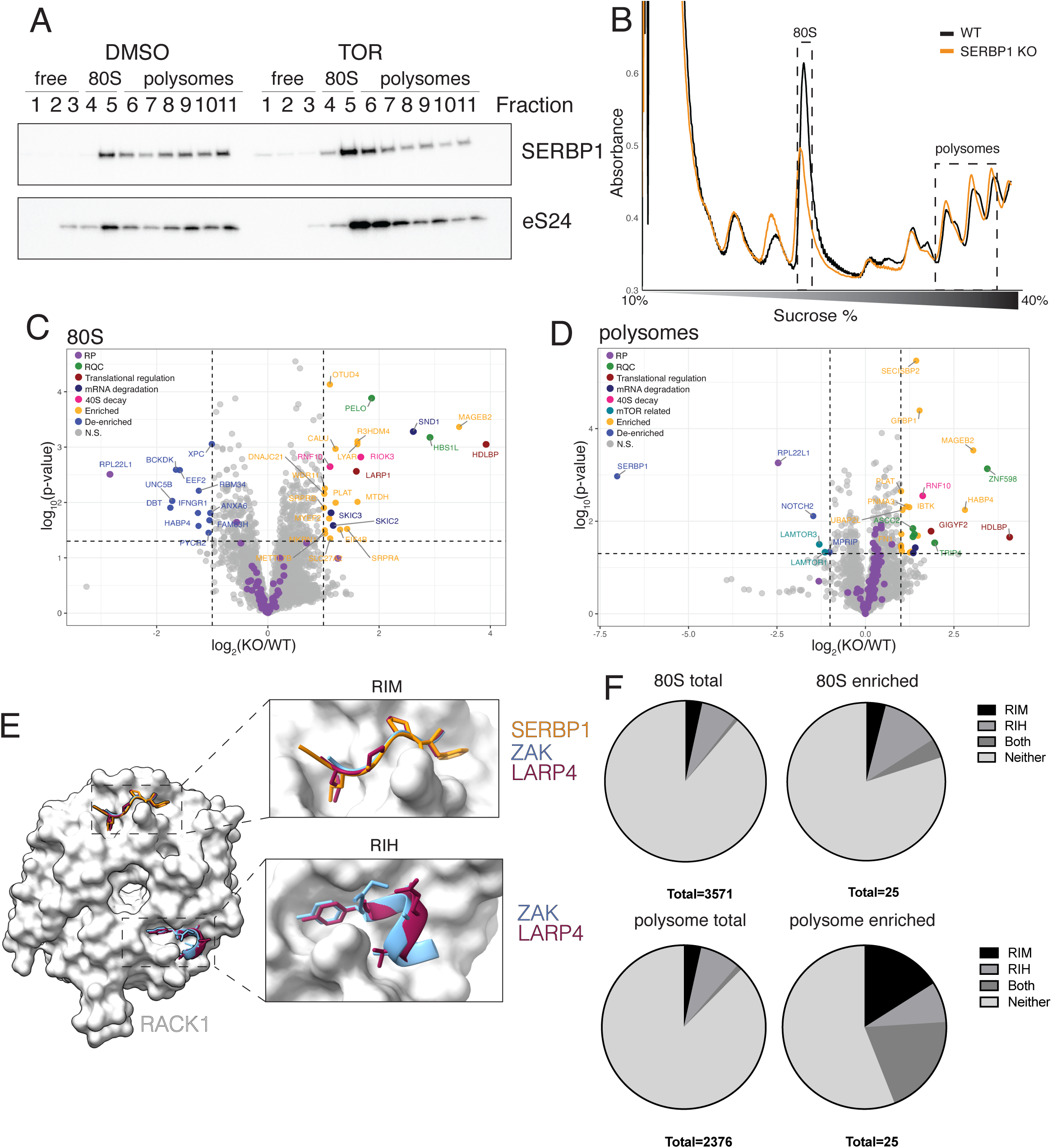
SERBP1 regulates the ribosome interactome. A) Western blot of gradient fractions from DMSO or torin treated cells using antibodies against SERBP1 or the ribosomal protein eS24. B) Absorbance traces from sucrose gradient sedimentation for WT or SERBP1 KO cells. Mass spectrometry was performed on the highlighted fractions containing monosomes or polysomes. C-D) Volcano plots for relative quantities obtained from mass spectrometry showing the log_2_(SERBP1 KO/WT) on the x-axis and the log_10_(p-value) on the y-axis for (C) 80S or (D) polysomes (n=4). Ribosomal proteins (RP) are shown in purple, samples with an enrichment of more than 2 fold and a p-value <0.05 are highlighted in orange, while samples that are de-enriched by more than 2 fold and have a p-value <0.05 are shown in blue. Proteins within relevant functional categories (ribosome quality control (RQC), translational regulation, mRNA decay, mTOR-related and 40S decay) are highlighted using different colors. E) Alphafold3 models for RIMs and RIHs from SERBP1, LARP4 and ZAK on RACK1. F) Pie charts showing percentages of proteins with RIMs, RIHs or both that are included in the 80S and polysome analyses shown in figures 1C and 1D or that are enriched on monosomes or polysomes upon SERBP1 depletion.

We next explored the underappreciated role of SERBP1 on elongating ribosomes. Using puromycin incorporation assays to monitor global translation, we first asked whether depletion of SERBP1 (using siRNAs or in a CRISPR-Cas9 knockout cell line) affected global translation in HEK-293T cells. Despite its presence on elongating ribosomes, the loss of SERBP1 had, at most, modest impacts on translational output as assessed by puromycin incorporation (Figure S1A-C). To more carefully examine the impact of SERBP1 on rates of translation elongation, harringtonine (HTN) run off assays were performed. Similarly, these assays revealed little to no differences in ribosome clearing rates (Figure S1D-E). Given the known role of SERBP1 in competing with the collision-responsive kinase, ZAKα,^10^ we checked if depletion of SERBP1 leads to an increase of basal collisions by examining the recruitment of the collision marker EDF1^23,24^ to polysomes in WT or SERBP1 KO cells that were either untreated or treated with collision-inducing doses of the elongation inhibitor anisomycin (ANS). As expected, EDF1 shifts to polysomes in WT cells treated with ANS but there was little to no EDF1 recruitment on polysomes in untreated WT or SERBP1 KO cells (Figure S1F). We conclude that under basal conditions, SERBP1 does not play a role in preventing collisions. Together these data suggest that SERBP1 binding to polysomes neither accelerates nor diminishes overall rates of translation.

### SERBP1 binding impacts the ribosome interactome

To gain further insight into possible roles of SERBP1 on polysomes, we asked how SERBP1 impacts the ribosome interactome by performing mass spectrometry on whole-cell lysates or on monosome- or polysome-containing sucrose gradient fractions from WT or SERBP1 KO cells (Figure 1B). Analysis of total lysates did not reveal major changes to the proteome upon SERBP1 depletion as most proteins do not experience an increase or decrease in levels greater than 2-fold (Figure S2A); the exception is MAGEB2 whose abundance is substantially increased in SERBP1 KO cells. Importantly, however, knockout of SERBP1 led to substantive changes in proteins associated with both monosomes and polysomes (Figures 1C-D).

On monosomes, SERBP1 depletion leads to an enrichment of factors known to associate with both translating and non-translating 80S, including Pelota, LARP1 and components of the SKI complex (Figure1C).^25–28^ These data also reveal an enrichment of two factors known to play a role in specifying 40S degradation: the E3 ubiquitin ligase RNF10 and the atypical kinase RIOK3.^17–21^ Finally, the dataset reveals previously unappreciated interactions between ribosomes and a number of factors including MAGEB2. As expected, loss of SERBP1 reduced interactions between ribosomes and SERBP1’s known binding partner eEF2 (Figure 1C).^2,3^

On polysomes, SERBP1 knockout led to an enrichment of the RQC factors ZNF598, ASCC1, ASCC2, and ASCC3 (Figure 1D), all of which are known to be recruited to collided ribosomes. Since SERBP1 depletion does not increase collisions (Figure S1F), we conclude that it instead prevents sporadic interactions between ribosomes and the RQC machinery. Importantly, SERBP1 depletion leads to an enrichment of many other factors on polysomes including RNF10, CUL3, GIGYF2, MAGEB2 and HDLBP, suggesting that it may play a broad role in preventing incidental interactions between ribosomes and these factors involved in various regulatory pathways. Together, these data reveal that SERBP1 impacts ribosome binding of many factors known to regulate diverse ribosome-centric functions.

SERBP1 has been shown to regulate the activity of the ZAKα kinase by competing with ZAK for a hydrophobic pocket found on the ribosomal protein RACK1.^10^ This competition was revealed in cryo-EM studies showing the presence of a so-called RACK1-interacting motif (RIM) on both SERBP1 and ZAK (Figure 1E);^10^ interestingly, a similar motif was also reported on another protein involved in RNA metabolism LARP4.^14^ Both LARP4 and ZAK also have a so-called RACK1-interacting helix (RIH) as defined in the same cryo-EM study, which was also shown to be critical for ZAK binding to ribosomes (Figure 1E).^10^ These previous observations suggested that RACK1 might be an important regulatory hub for ribosome interactors.

To further explore the importance of the SERBP1-RACK1 interaction, we looked for putative RIM and RIH motifs in proteins that were significantly enriched in our mass spectrometry datasets. Putative RIMs were identified on 4.5% and 6% of proteins detected on monosomes or polysomes, respectively, and on 8% and 20% of proteins enriched on monosomes and polysomes in SERBP1 KO cells. Similarly, putative RIHs were found on 8% and 9% of proteins detected on monosomes or polysomes, respectively, and in 16% and 28% of proteins enriched in these fractions upon SERBP1 KO (Figure 1F; Table S1). These observations suggest that the site of interaction between RACK1 and SERBP1, ZAK and LARP4 (the RIM) as well as the second common site of RACK1 binding for ZAK and LARP4 (the RIH) may be broadly involved in ribosome-mediated stress responses, and that SERBP1 may thus play an extensive regulatory role.

### Stress conditions reveal additional ribosome interactions regulated by SERBP1

Since depletion of SERBP1 gave rise to many interactions between ribosomes and proteins involved in stress responses (eg. RQC components), we wanted to further explore the impact of SERBP1 in regulating interactions in the context of stress. To do so, we performed mass spectrometry on monosomes and polysomes from WT and SERBP1 KO cells treated with two drugs that impact translation in different ways: torin-1 (TOR), which inhibits mTOR leading to a widespread decrease in translation initiation, and anisomycin (ANS), which induces collisions and leads to the activation of the ribotoxic stress response (Figure 2A). While these drug treatments generally led to modest changes in the total levels of certain proteins in the absence of SERBP1 (Figure S2A), these data also revealed proteins that were differentially enriched with ribosomes upon SERBP1 depletion in the context of TOR or upon ANS treatment (Figures 2B-C). ANS treatment in SERBP1 KO cells resulted in an enrichment of several subunits of the CCR4-NOT complex on ribosomes (Figure 2B-C, right) consistent with previous studies indicating connections between ribosome collisions and mRNA decay pathways.^29,30^ Similarly, TOR-treatment revealed additional factors involved in mRNA degradation, translational regulation and ribosome quality control upon SERBP1 depletion (Figure 2B-C, left) including exosome and CCR4-NOT components, EDF1 and GIGYF2. Interestingly, SERBP1 depletion in TOR-treated cells and in unstressed cells led to a de-enrichment of several factors on polysomes including the “ragulator” components, LAMTOR1 and LAMTOR3 (Figures 1D and 2C), which could reflect direct interactions between the ribosome and these critical mTOR regulators. We also identified a group of proteins that were enriched on ribosomes in all three conditions: untreated, ANS- or TOR-treated (Figure 2D-E). These included RQC factors, mRNA decay machinery, as well as RIOK3 and RNF10, both previously implicated in selective 40S degradation. Our datasets reveal a broad regulatory landscape for SERBP1 in interacting with the ribosome in stressed and unstressed cells.

**Figure 2:**
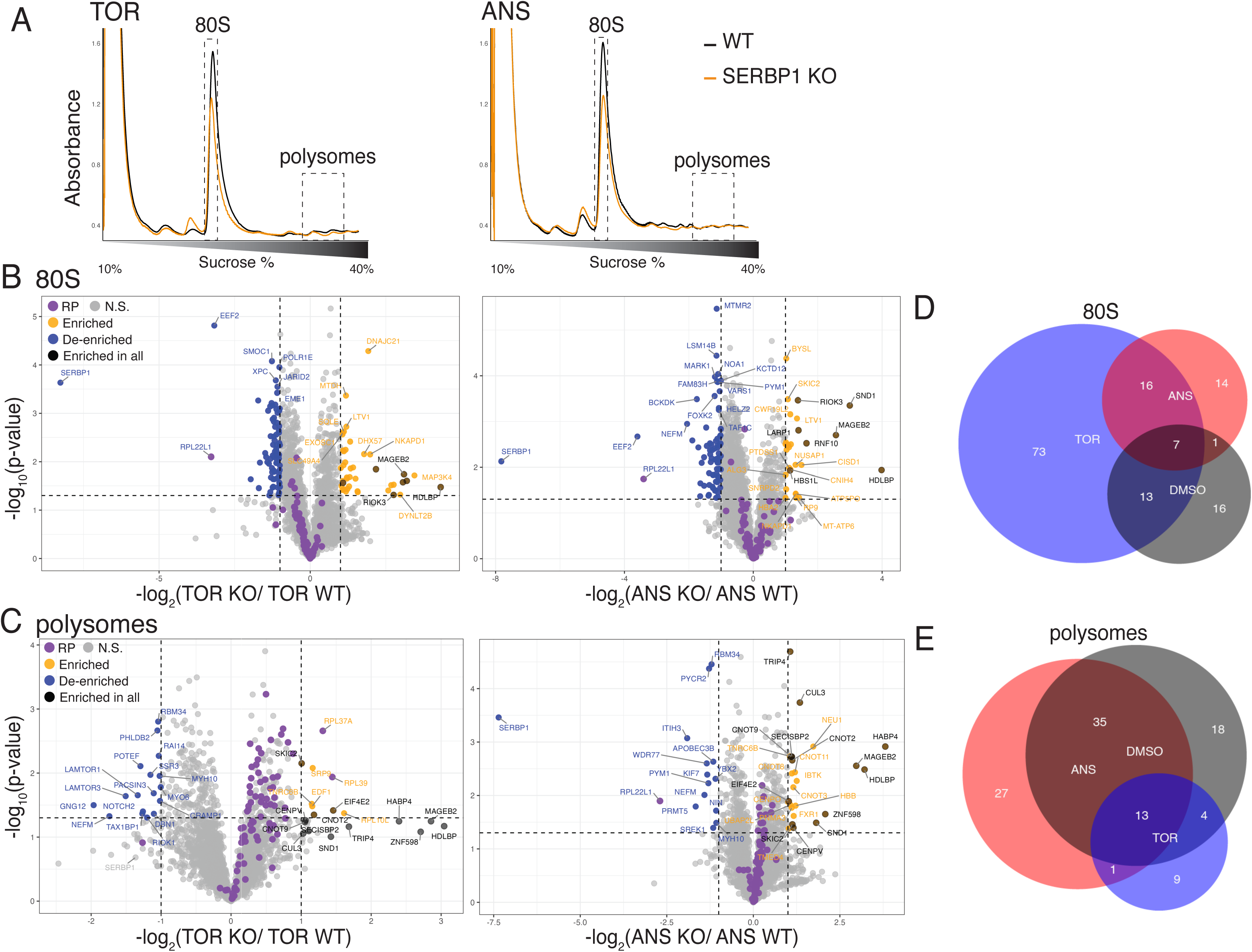
Stress conditions highlight protective roles of SERBP1 on ribosomes. A) Absorbance traces from sucrose gradient sedimentation for WT or SERBP1 KO cells treated with torin-1 (TOR) or anisomycin (ANS). B-C) Volcano plots for relative quantities obtained from mass spectrometry showing the log_2_(SERBP1 KO/WT) on the x-axis and the log_10_(p-value) on the y-axis for (B) 80S or (C) polysomes from TOR or ANS treated cells (n=4). Ribosomal proteins (RP) are shown in purple, samples with an enrichment of more than 2 fold and a p-value <0.05 are highlighted in orange, while samples that are de-enriched by more than 2 fold and have a p-value <0.05 are shown in blue. Proteins enriched in DMSO, TOR and ANS-treated samples are shown in black. D-E) Venn diagrams showing the overlap of factors enriched by a fold change of more than 2 after DMSO, ANS and/or TOR treatment.

### SERBP1 protects 40S subunits from stress-induced degradation

As a proof of principle, we focused on the impact of SERBP1 depletion on ribosome homeostasis because of the observed SERBP1-dependent enrichment of RNF10 and RIOK3 on ribosomes across all conditions. Additionally, it has already been documented in the literature that SERBP1 plays a role in protecting ribosomes from degradation upon mTOR inhibition.^5^ To test the impact of SERBP1 on ribosome distribution, we started by depleting SERBP1 and treating cells with DMSO or TOR for 16 hours. Consistent with the widespread translation inhibition caused by TOR, we observe an increase in monosomes accompanied by a corresponding decrease of polysomes in TOR-treated cells compared to DMSO-treated cells (Figure 3A, blue and black traces). We saw similar decreases in polysomes upon treatment with torin in cells where SERBP1 is knocked down with siRNAs. However, SERBP1 depletion by knockdown or knockout led to a distinctive decrease in the monosome peak and an increase in the free 60S subunit peak when compared to WT cells treated with TOR (Figures 3A and S3A orange and blue traces). The increase in 60S subunits is reminiscent of earlier reports where an increase in 60S subunits was shown to reflect a relative loss of 40S subunits caused by RIOK3/RNF10-mediated degradation.^17,20^

**Figure 3:**
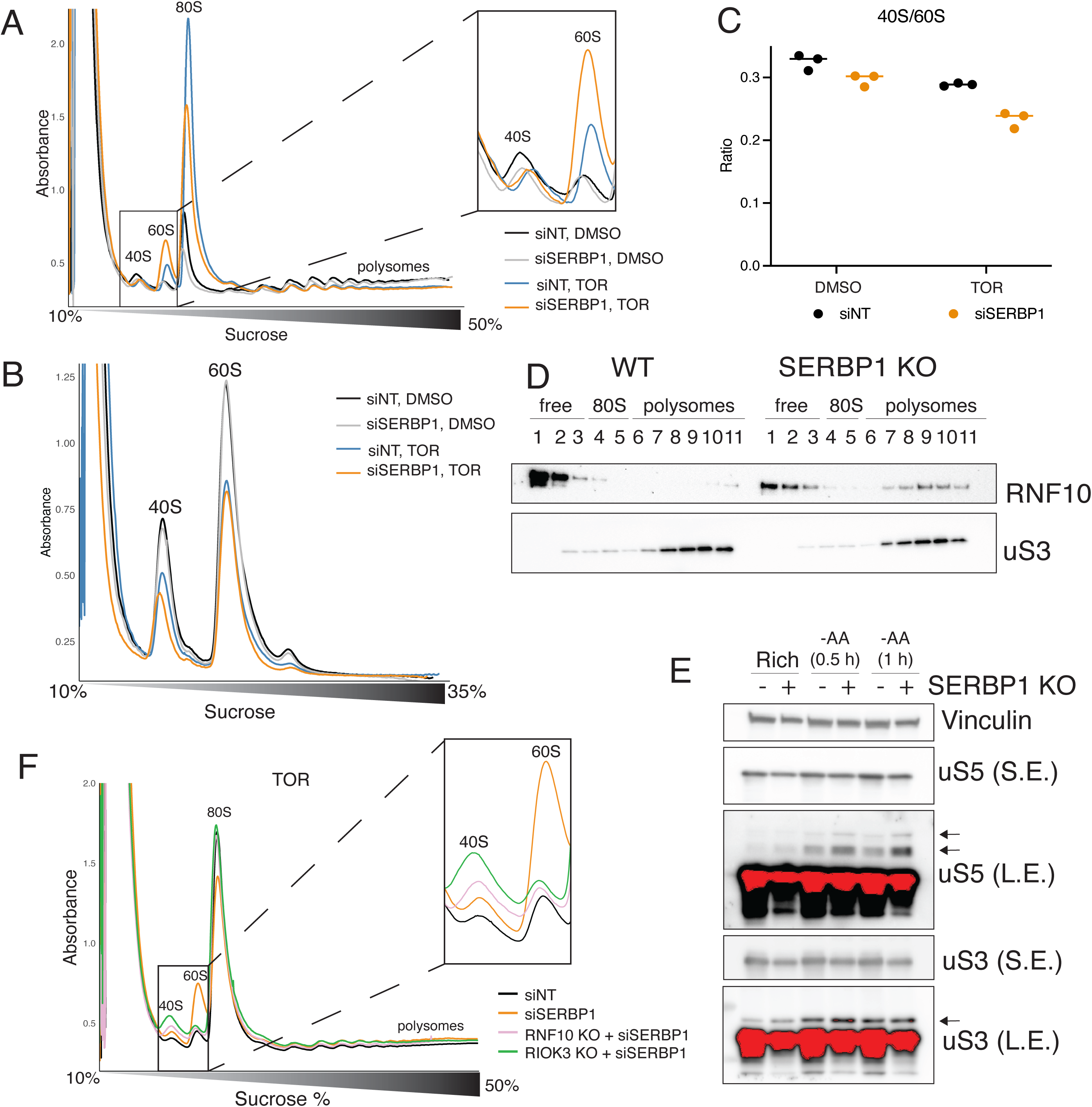
SERBP1 depletion induces selective 40S degradation. A) Absorbance traces from sucrose gradient sedimentation of samples that were treated with DMSO or torin and transfected with non-targeting siRNAs (NT) or siRNAs against SERBP1. A closer view of the 40S and 60S peaks is highlighted. B) Absorbance traces from sucrose gradients of EDTA-split lysates from untreated or torin-treated cells transfected with NT siRNAs or siSERBP1. Traces were aligned on the x-axis to facilitate comparisons. C) Quantification of the area under the curve observed from the 40S and 60S peaks for cells transfected with NT siRNAs or siSERBP1 shown in B. D) Western blot of gradient fractions from WT or SERBP1 KO cells using an antibody against RNF10 or the ribosomal protein uS3. E) Western blots of WT or SERBP1 KO cells kept in rich media or exposed to amino acid starvation for 0.5 or 1 hour using an antibody against the loading control vinculin or against the ribosomal proteins uS5 or uS3 (S.E. = short exposure; L.E.= long exposure). Arrows highlight ubiquitylated species. F) Absorbance traces from sucrose gradient sedimentation of torin-treated WT cells transfected with NT siRNAs and WT, RNF10 KO or RIOK3 KO cells transfected with siSERBP1. A closer view of the 40S and 60S peaks is highlighted.

To test whether the increase in 60S subunits reflects decreased levels of 40S subunits upon SERBP1 knockdown, we quantified the total abundance of each subunit by dissociating all 80S ribosomes into 40S and 60S subunits and performing sucrose gradient sedimentation after normalizing the lysates to cell count (Figure 3B). While torin treatment resulted in a clear loss of both subunits as previously reported,^5^ quantitation of the 40S and 60S peaks revealed a modest decrease in 40S subunits relative to 60S both as a function of torin treatment and as a function of SERBP1 depletion. Importantly, the decrease in the 40S/60S ratio is exacerbated in torin-treated cells where SERBP1 was depleted (Figures 3B-C). These data confirm that the observed increase in the 60S peak reflects a selective decrease in 40S subunits and represents a convenient proxy to follow this process. Altogether, these data suggest that SERBP1 plays a critical role in protecting ribosomes from degradation basally, and this protective role becomes even more apparent under conditions of mTOR inhibition.

We next wanted to directly test the involvement of our mass spectrometry hits, RNF10 and RIOK3, on the observed loss of 40S subunits as they have both been previously implicated in a pathway that mediates the selective degradation of 40S subunits.^17–19^ We validated our mass spectrometry observation that RNF10 is enriched on polysomes in SERBP1 KO cells by performing western blots on sucrose gradient fractions from WT and KO cells (Figure 3D). Next, we evaluated the impact of SERBP1 on RNF10 ubiquitylase activity by exposing WT or KO cells to amino acid starvation, a potent activator of RNF10.^17^ Indeed, we find that the levels of ubiquitylation of RNF10 targets uS5 and uS3 increase in SERBP1 KO cells compared to WT after 30 or 60 minutes of starvation (Figure 3E). We then asked if RNF10 and the downstream effector RIOK3 were responsible for the observed 40S degradation in SERBP1-depleted cells by co-depletion of SERBP1 in either RNF10 or RIOK3 KO cells (Figure S3B). Sucrose gradient traces obtained from these torin-treated cells reveal that depleting either RNF10 or RIOK3 in combination with SERBP1 blocks the increase in the 60S peak that reflects 40S degradation observed after depletion of SERBP1 alone (Figure 3F).

Our mass spectrometry data also revealed an enrichment of components of the ASC-1 complex on polysomes in the absence of SERBP1 (Figure 1D & Table S1). In particular, ASCC3 has been shown to work downstream of RNF10 to split ribosomes and release 40S subunits for degradation during starvation.^17^ Indeed, depletion of ASCC3 in SERBP1 KO cells also leads to a decrease in the 60S peak reflecting the blocking of 40S degradation (Figure S3C). Together, these data demonstrate that SERBP1 protects 40S subunits from degradation mediated by RNF10, RIOK3 and ASCC3, and all three factors were revealed to be enriched in the absence of SERBP1 in our mass spectrometry experiment.

### SERBP1 competes with RIOK3 to protect 40S subunits from degradation

Given that SERBP1 has multiple binding sites (or handles) on the ribosome, we wanted to test which of these sites of interactions with the ribosome were critical for protection against RNF10/RIOK3 mediated degradation. We hypothesized that RIOK3 might compete with SERBP1 at the mRNA binding channel of the ribosome because both of these factors are known to bind there (Figure 4A).^17^ To test if the region of SERBP1 that binds the mRNA channel is important for preventing 40S degradation, we generated a SERBP1 deletion mutant (ΔmRNA channel binding; Figure 4B) and first made certain that this deletion mutant was well expressed (Figure S4A). We observed that the ΔmRNA channel binding SERBP1 is still able to interact with ribosomes presumably through its other binding handles on the 60S subunit and the 40S RACK1 binding site (Figure S4B). We then transfected full length or ΔmRNA channel binding constructs into TOR-treated SERBP1 KO cells and performed sucrose gradient sedimentation on these lysates. Cells transfected with full length constructs show a decreased 60S peak compared to KO cells, consistent with a restoration of SERBP1 function (Figure 4C); in contrast, in cells transfected with the ΔmRNA channel binding construct, we see an intermediate phenotype consistent with a partial but important contribution of SERBP1 to blocking RIOK3 function through the mRNA channel site (Figure 4C). Since deletion of the mRNA channel binding region of SERBP1 may have unanticipated effects on how SERBP1 interacts with 40S subunits, we also generated SERBP1 mutants where the positively charged amino acids in the region that interact with the mRNA binding channel were mutated to either aspartic acid (reverse charge) or alanine (neutral charge). Consistent with the results obtained from the ΔmRNA channel construct, SERBP1 mutants with reversed or neutralized charges in the region that binds the mRNA channel also show an intermediate rescue of the torin-induced 40S degradation phenotype (Figure S4C). Together, these results argue that the mRNA channel binding region of SERBP1 is important for 40S protection from RIOK3.

**Figure 4:**
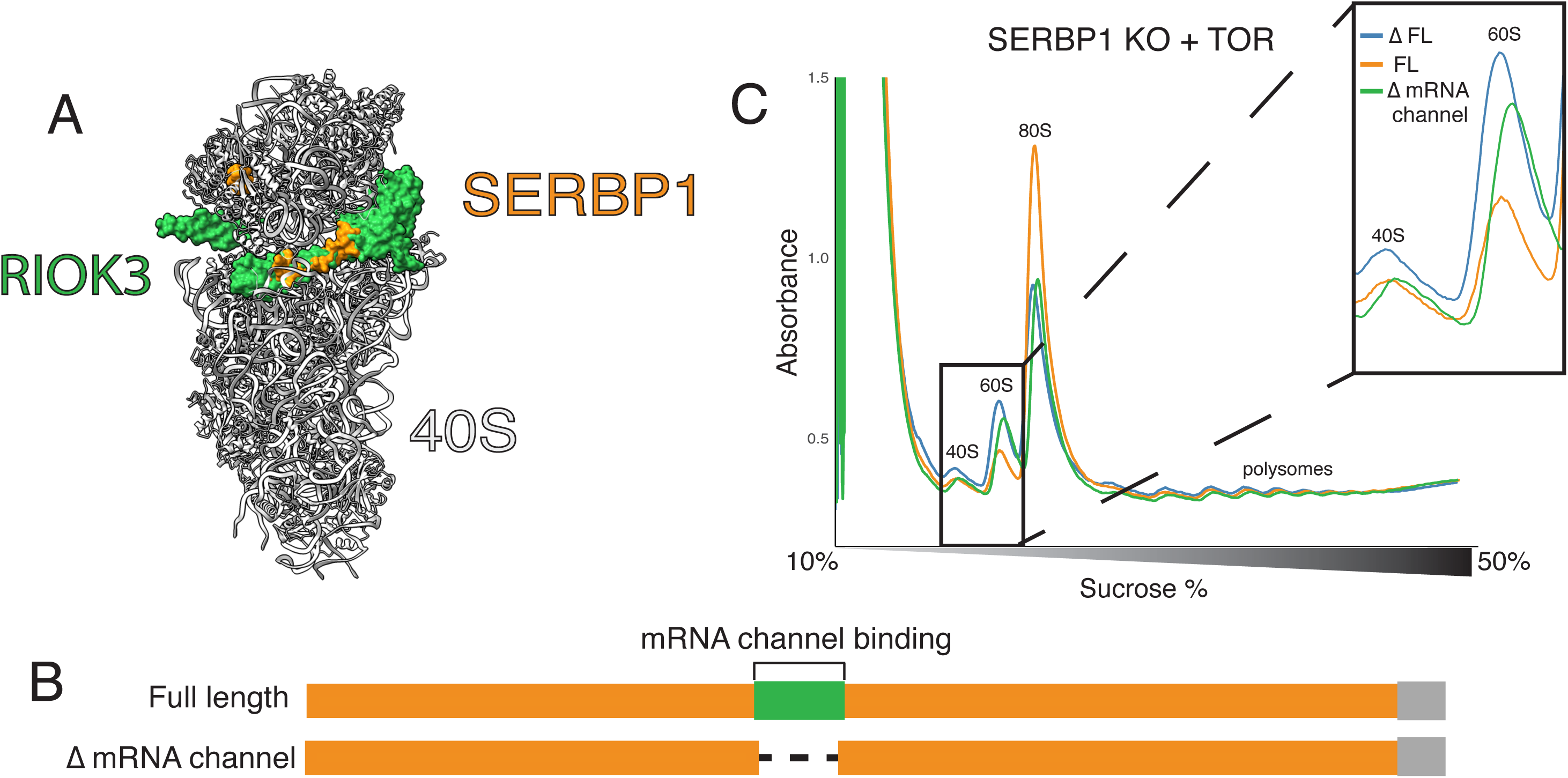
SERBP1 competes with RIOK3 to protect ribosomes from degradation. A) Overlay of the structure of RIOK3 (PDB: 8ZDB) and the structure of SERBP1 (PDB: 6Z6M) on the 40S subunit. B) Schematic representation of SERBP1 constructs. C) Absorbance traces from sucrose gradient sedimentation of torin-treated SERBP1 KO cells transfected with plasmids expressing eGFP as a control, full length SERBP1 or a SERBP1 truncation lacking the region binding the mRNA channel.

### RNF10 motif predicted to overlap with SERBP1 on RACK1 regulates ubiquitylation

Our mass spectrometry data show an enrichment of RNF10 on both monosomes and polysomes upon SERBP1 depletion (Figures 1C-D & 3D), and, as previously mentioned, the C-terminus of SERBP1 interacts with RACK1, which is next to one of RNF10’s targets, uS3, on the ribosome. Moreover, *in vitro* studies measuring uS3 ubiquitylation have shown that RACK1’s yeast homologue (Asc1) is important for the activity of RNF10’s yeast homologue Mag2.^31^ To test if RACK1 might directly interact with RNF10, we explored interactions using AlphaFold3. These predictions reveal two potential sites of interaction between RNF10 and RACK1: a putative RIM and a putative RIH (Figure 5A & S5A-B). These predictions suggest there might be simple steric conflicts between RNF10 and SERBP1’s own RIM.

**Figure 5:**
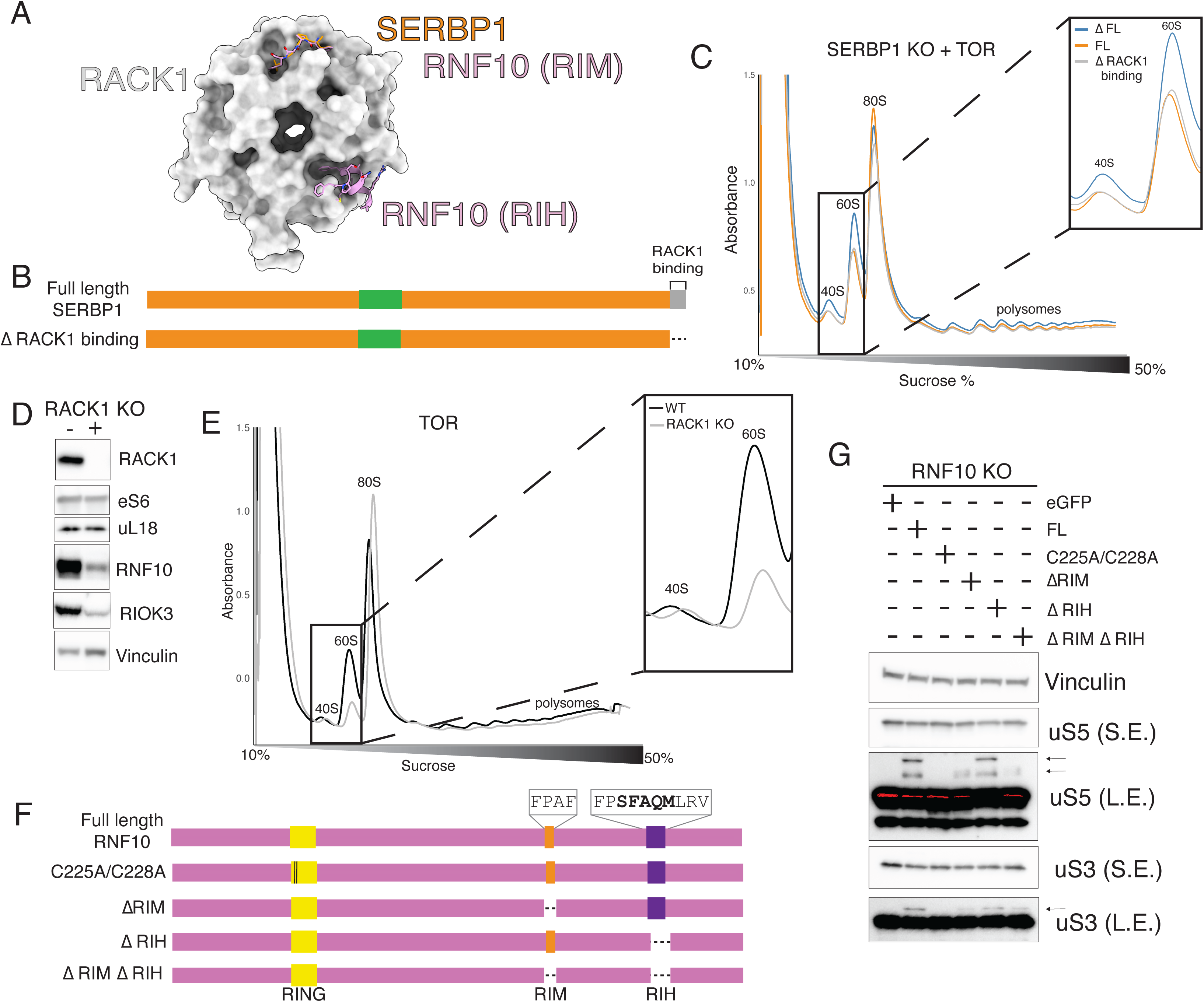
RNF10 RIM promotes ubiquitylation of uS3 and uS5. A) Overlay of the structure of SERBP1 (PDB: 9RPV) and the alphafold prediction of RNF10 on the ribosomal protein RACK1. B) Schematic representation of SERBP1 constructs. C) Absorbance traces from sucrose gradient sedimentation of torin-treated SERBP1 KO cells transfected with plasmids expressing eGFP as a control, full length SERBP1 or a SERBP1 truncation lacking RACK1 binding region. D) Western blots from WT or RACK1 KO cells using antibodies against vinculin (loading control), RNF10, RIOK3, eS6 or uL18. E) Absorbance traces from sucrose gradient sedimentation of torin-treated WT or RACK1 KO cells transfected with plasmids expressing RNF10 and RIOK3. F) Schematic representation of RNF10 mutants and truncations. Deleted sequences are highlighted in a box above the respective regions. The five conserved residues of the RIH are shown in bold. G) Western blots from WT or RNF10 KO cells using antibodies against vinculin (loading control), uS5 or uS3 (S.E. = short exposure; L.E.= long exposure). Ubiquitylated species are shown with arrows.

To test this model, we created a construct expressing a truncated version of SERBP1 lacking the region that interacts with RACK1 (ΔRACK1 binding; Figure 5B). This truncation mutant expresses well and still interacts with elongating ribosomes (Figure S5C-D), likely through the other known interaction site on the 60S and 40S. Sucrose gradient sedimentation experiments on torin-treated cells expressing this ΔRACK1 SERBP1 binding construct revealed that both full length SERBP1 and the ΔRACK1 binding mutant equivalently diminish the 60S peak (and thus protect the 40S subunit from degradation) (Figure 5C). Therefore, this region of SERBP1, predicted to compete with RNF10 for RACK1 binding, does not on its own impact 40S protection. Since the ΔRACK1 binding SERBP1 still effectively binds to 40S subunits (Figure S5D) and likely maintains interactions at the mRNA binding channel that would compete with RIOK3, we were not surprised to see that the deletion mutant failed to reveal a phenotype in 40S degradation.

While the RACK1 binding region of SERBP1 is dispensable for protection of 40S from degradation, we wondered if RACK1 itself was important for this process. As RACK1 has been previously shown to be essential for uS3 ubiquitylation,^32^ we wanted to deplete RACK1 to investigate its role in RNF10/RIOK3 mediated degradation. However, decreasing 40S levels leads to lower levels of RNF10.^17,33^ To test if the targeted knockdown of RACK1 levels was sufficient to reduce RNF10 levels, we looked at RNF10 levels in RACK1 knockout cells and in WT cells where RACK1 was knocked down. Indeed, our data show that RNF10 and RIOK3 levels do substantially decrease in the absence of RACK1 (Figures 5D & S5E), thus providing a potential explanation for previous studies indicating that RACK1 is essential for uS3 ubiquitylation.^32^

To preserve the degradation pathway components RNF10 and RIOK3 in RACK1 KO cells, we overexpressed both proteins in WT and RACK1 KO cells (Figure S5F) and then asked whether RACK1 was required for degradation in TOR-treated cells. Sucrose gradient sedimentation of these samples reveals that knocking out RACK1 protein disrupts 40S degradation even in the presence of abundant levels of RNF10 and RIOK3 (Figure 5E). These data argue that RACK1 is indeed required for promoting RNF10/RIOK3-mediated 40S degradation.

In light of this, we decided to test if the RNF10 motifs predicted to interact with RACK1 are important for RNF10-mediated ubiquitylation. We prepared a series of mutants including deletion of the RIM (ΔRIM), the RIH (ΔRIH), or both motifs (ΔRIM ΔRIH), as well as a catalytically dead (C225A/C228A) variant (Figure 5F & Figure S5G) and expressed them in cells deficient for RNF10. Western blots for uS3 and uS5 in experiments where these variants were transfected into cells show that full length (FL) RNF10 is sufficient to induce ubiquitylation compared to either KO cells transfected with an eGFP control or cells transfected with the C225A/C228A construct (Figure 5G). By contrast, the ΔRIM RNF10 variant showed impaired ubiquitylation of both uS3 and uS5, while the ΔRIH variant was unimpaired relative to FL. Cells transfected with the construct lacking both motifs showed ubiquitylation levels similar to that of the ΔRIM variant (Figure 5G). We conclude that the RIM is critical for RNF10 activity, while the RIH is dispensable.

We next asked whether the two RACK1-interaction motifs on RNF10 are conserved among other species and further compared their sequences to other proteins known to bind these RACK1 sites. The RACK1-interacting motif of RNF10 is evolutionarily well-conserved (Figure S5I) and, as previously mentioned, is similar to the FPxL motif of SERBP1 and ZAK as well as the FPxL motif on LARP4 (Figure S5B & H).^10,14^ Interestingly, there may be some variation on these conserved sequences that is becoming apparent with the expansion of the set. For example, on RNF10, the last amino acid of this tetrad is a phenylalanine instead of a leucine (Figures S5H-I), suggesting that this site on RACK1 may simply favor binding by nonpolar amino acids, but not specifically leucine (FPXΦ). Expanding our analysis of proteins enriched for putative RIMs to now include both FPxL and FPxF reveals that, while these account for only 6% of all proteins identified on ribosomes in the mass spectrometry analyses, proteins containing a putative RIM account for 16% and 24% of the proteins enriched on monosomes and polysomes, respectively, upon SERBP1 depletion (Figure 1F, Table S1). The second predicted RACK1-interacting motif of RNF10 is composed of five relatively well-conserved amino acids and again resembles ZAK and LARP4 sequences known to bind RACK1 at this second site. Interestingly, we also find a putative RIH motif on USP10, which removes the ubiquitins from uS3 and uS5, and find that it is also well conserved (Figures S5B and S5H). While these five amino acids demonstrate variability between the proteins evaluated, they follow a pattern where the five amino acids are composed of a serine followed by an aromatic amino acid (Ω), an alanine, any amino acid and a hydrophobic amino acid (Φ) (SΩAXΦ) (Figure S5H-I). These data highlight two different well-conserved motifs that are prevalent amongst proteins that interact with ribosomes and are significantly enriched on ribosomes in the absence of SERBP1.

## Discussion

Here, we show that SERBP1 serves protective roles on ribosomes preventing interactions between them and factors involved in different pathways, including ribosome quality control, translational regulation, RNA decay and ribosome degradation. As a case study, we demonstrate that SERBP1 protects 40S subunits from degradation by RNF10 and RIOK3. We propose that this occurs through a two-step mechanism. In the first step, SERBP1 binding to key sites on RACK1 on the 40S blocks RNF10-mediated ubiquitylation. Then, in a second step, even if this initial protection is bypassed, SERBP1 is still protective through its interactions with the mRNA channel of the 40S subunit that blocks RIOK3 binding (Figure 6). This work expands our understanding of the role that SERBP1 plays in regulating ribosome homeostasis. Additionally, our deep proteomic data suggest that SERBP1 plays much broader protective roles in stressed and unstressed cells.

**Figure 6:**
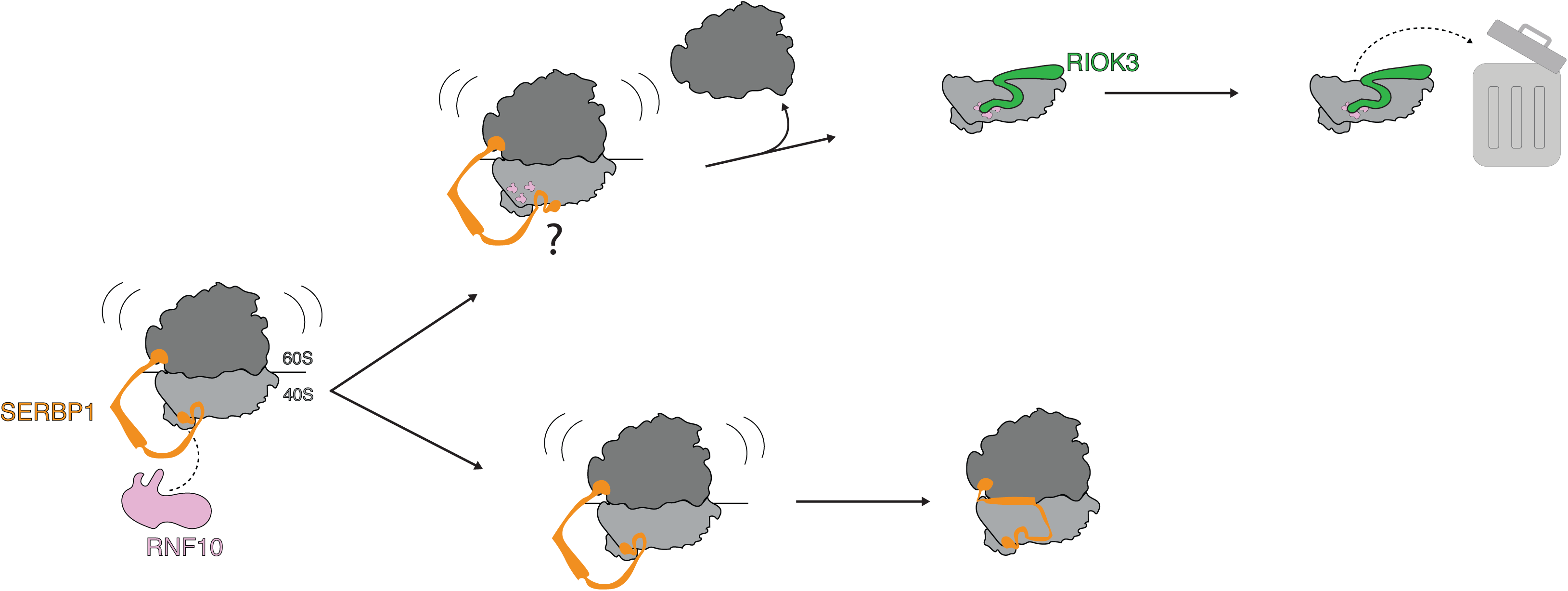
SERBP1 provides a two-step layer of protection from selective degradation of 40S subunits. Cartoon representation of our model showing SERBP1 protecting ribosomes from ubiquitylation by RNF10. If this protection is bypassed, ribosomes can still be protected by SERBP1 through its binding of the mRNA channel of the ribosome where it obstructs recognition of the ubiquitylated sites by RIOK3.

SERBP1 and its yeast homolog Stm1 have documented roles in bulk ribosome degradation as well as in biogenesis, with some studies reporting that tagged versions of SERBP1 localize to the nucleolus.^5,34,35^ We cannot discard the possibility that a defect in biogenesis contributes to the global decrease in both subunits observed in cells depleted of SERBP1 upon torin treatment. However, a decrease in early biogenesis seems unlikely to play a role in the selective loss of 40S subunits that we observe, especially given that our phenotype is dependent on RNF10 and ASCC3, both of which have well established roles in cytoplasmic-driven quality control.^36–38^ Importantly, however, RIOK3 has been observed to bind to cytosolic late stage biogenesis intermediates, potentially confounding a simple interpretation.^17^

SERBP1 binding to ribosomes has long been regarded as a hallmark of dormancy.^2,3,39^ However, there is increasing evidence that SERBP1 binds polysomes as well as monosomes, taking advantage of binding handles located on portions of the ribosome distinct from the mRNA binding channel.^7–9^ These observations are compatible with the fact that SERBP1 is extremely abundant, present almost at a 1:1 stoichiometry with ribosomes in human cells, and is abundantly found on elongating ribosomes,^6^ possibly facilitating the transition between dormant and elongating states. We detect an increased association of RNF10 in both the monosomes and polysomes of cells depleted of SERBP1 even when cells are unstressed.

Therefore, we speculate that SERBP1’s association with both monosomes and polysomes plays important roles in maintaining ribosome levels by blocking sporadic binding of RNF10 to ribosomes even during unstressed conditions and preventing incidental targeting of 40S subunits for degradation.

Our mass spectrometry data reveal a wealth of ribosome interactors that are antagonized by the presence of SERBP1 and the recently described structural handles for SERBP1 binding on the ribosome rationalize where such competition can occur. Our work highlights the importance of the regions of SERBP1 that bind to the mRNA channel of the ribosome in competing with RIOK3 to prevent 40S degradation. LARP1, another protein known to bind the mRNA channel of the ribosome,^28,40^ was similarly enriched in monosomes obtained from SERBP1 KO cells in stressed and unstressed cells. Interestingly, when LARP1 is found in its “TOP complex” – bound to 40S subunits and to TOP mRNAs encoding ribosomal proteins – it inhibits their translation.^41^ As such, SERBP1 binding to the mRNA channel would compete with LARP1 binding and thus could decrease translational repression of the TOP mRNAs, feeding into more global ribosome homeostasis.

Interestingly, upon anisomycin treatment, we observe an enrichment of proteins related to mRNA degradation on ribosomes, the CCR4-NOT complex. This enrichment is consistent with the literature that reports an increase in mRNA decay associated with ribosome collisions in yeast and bacteria.^29,30^ The CCR4-NOT complex is well known to be involved in co-translational mRNA decay and indeed, several subunits of the CCR4-NOT complex have predicted RIMs (CNOT1, CNOT2 & CNOT4), suggesting a mechanism by which SERBP1 might regulate co-translational mRNA decay. Moreover, TOR-treatment leads to the enrichment of additional factors involved in mRNA decay, including components of the exosome, further implicating SERBP1 as a regulator of mRNA stability.

RACK1 clearly serves as a scaffold for many proteins to interact with ribosomes. Here, we find that RNF10, which has an FPxF motif that closely resembles the RIM in ZAK and SERBP1, is similarly predicted to bind RACK1. We demonstrate that this motif is important for RNF10 activity. Together with the observation that SERBP1 depletion enhances RNF10 activity, we provide evidence for a second regulatory process that is antagonized by the binding of SERBP1 to RACK1. Moreover, we find that 20-24% of proteins whose interaction with ribosomes is enriched in the absence of SERBP1 contain a putative RIM (Table S1), suggesting that this RACK1 binding site is a common place on the ribosome for regulatory factors to bind. Further study of the criteria required to prioritize binding by one factor versus the other to this site will provide crucial insight into how ribosomes use this platform to regulate stress responses. Interestingly, RACK1 is an unusual eukaryotic-specific ribosomal protein that may have arisen to play a more nuanced role in higher order ribosome function.

## Resource availability

Reagents generated for this study are available upon request, which will be fulfilled by the lead contact, Rachel Green.

## Materials and Methods

### Cell culture

HEK-293T cells (ATCC) were cultured in DMEM (10% FBS) and kept in a humidified incubator at 37°C at 5% CO_2_. For all experiments, RACK1 and SERBP1 KO cell lines were seeded at 1.5x the density of WT HEK-293Ts because they grow slower. For starvation experiments, cells were rinsed once with 5 mL of 1x PBS and then incubated in DMEM (10% dialyzed FBS) lacking amino acids. For cells kept in rich media, amino acids were added back to the media (DMEM, 10% dialyzed FBS). Torin treatments were performed by adding the drug to a concentration of 250 nM at least 1.5 hours after changing media and then incubating the cells with the drug for 16 hours or for 1 hour (mass spectrometry experiments). Anisomycin treatments were performed by treating cells with the collision-inducing dose of 0.2 µM for 1 hour. Puromycin incorporation assays were performed by incubating cells with 10 µg/mL of puromycin at 37°C for 15 minutes. Harringtonine run off assays were performed by adding 13 µM of harringtonine to the media and incubating cells for the indicated periods of time.

### siRNA knockdowns and plasmid transfections

For siRNA treatments or plasmid transfections, cells were seeded onto 10 cm^2^ plates at a density of 2×10^6^ for 24 h experiments, 1×10^6^ for 48 h experiments, 0.5×10^6^ for 72 h experiments or 0.3×10^6^ for 96 h experiments. siRNA knockdowns were performed using 50 nM SMARTpool siRNAs (Horizon discovery) targeting the transcript of interest and Lipofectamine RNAimax (Invitrogen) according to manufacturer’s instructions the day after plating the cells. Plasmids were transfected into cells using Lipofectamine 3000 according to manufacturer’s instructions 24 or 48 hours before harvesting.

### Cloning and plasmids

A gene block encoding SERBP1 was ordered from IDT. This was cloned into a modified pFC14K plasmid using gibson assembly. To match endogenous expression of SERBP1, the CMV promoter was swapped out for an eEF1a promoter. This construct was further modified to create SERBP1 truncations using PCRs to linearize the vector followed by ligation using KLD mixture (NEB). The plasmid for expression of RNF10 was a gift from the lab of Roland Beckmann. RNF10 is on a pcDNA5 plasmid under a full CMV promoter. Constructs expressing a 3x FLAG tag and a 2x Strep tag on the N or C terminus of RNF10 were tested. Mutants were generated by linearizing the plasmid and performing ligations using KLD. All plasmids were verified through whole plasmid sequencing (Plasmidsaurus) and are available upon request.

### Generation of CRISPR knock outs

Guides against RNF10 (GCAGCCAAGATAACCCGTTG) or SERBP1 (CACCACGACCTCGAATAGGT) were ordered from IDT and annealed using T4 PNK in T4 DNA ligation buffer at 37°C for 30 min followed by 5 min at 95°C and finalized by ramping down 5°C per min until reaching 25°C as indicated in Shalem *et al.* 2014.^42^ Annealed oligos were ligated at room temperature for 40 minutes using T4 ligase in T4 DNA ligation buffer onto a LENTI CRISPRv2 plasmid that had been digested using BsmBI and gel purified. Plasmids were then transformed into competent DH5 alpha cells that were then cultured at 30°C in carbenicillin resistant plates. Single colonies were picked and midiprep cultures were grown at 30°C. Midipreps were performed using the ZymoPURE II Plasmid Midiprep Kit. All plasmids were verified through whole plasmid sequencing.

For transfections, low passage (4 to 5) HEK-293T cells were seeded at 0.15×10^6^ cells onto a 6-well dish. The next day, cells were transfected as described above using 1.5 µg of plasmid. After 48 h, cells were trypsinized, passed through a cell strainer and diluted into single cells onto a 96-well plate. Once single colonies reached confluency, the cells were transferred to larger plates. After 2 to 3 passages, the expression of RNF10 or SERBP1 was verified through western blots and the respective genomic locus was verified through sequencing.

### Sucrose gradient sedimentation

For normal gradients, samples were rinsed once with 4mL of 1x PBS (360 µM emetine) and lysed in 200 µL of lysis buffer (50mM HEPES pH 7.4, 100mM KOAc, 5% Glycerol, 0.5% Triton X, 15mM Mg(OAc)_2_, 360µM emetine, 1x Halt Protease and Phosphatase Inhibitor, 80 units of Turbo DNase, 1mM TCEP). After scraping the samples, lysates were transferred to a 1.5 mL tube. Lysates were clarified by spinning at 8,000 xg at 4°C for 5 min. Samples were then flash frozen and stored at −80°C. Sucrose gradients were prepared by layering 6 mL of 50% or 40% sucrose buffer onto 6 mL of 10% sucrose buffer containing 1x gradient buffer (250 mM HEPES pH 7.5, 1M KOAc, 50mM MgOAc) into an open-top polyclear centrifuge tube (Seton). 10-40% (for mass spectrometry) or 10-50% gradients were made on a gradient master (Biocomp). Samples were normalized to RNA concentrations using Qubit BR RNA kit. At least 20 µg of RNA were loaded onto sucrose gradients in 350 µL. After balancing samples, they were spun for 1 h and 45 min or 2 h (for mass spectrometry) at 4°C, 40,000 rpm in a SW-41 rotor. Fractions were collected using a Biocomp Piston Gradient Fractionator that allows UV absorbance (260 nm) measurements to obtain traces.

For split gradients, samples were rinsed once with 4mL of cold 1x PBS and lysed in 150 µL of lysis buffer (20 mM Tris-Cl pH 7.5, 140 mM KCl, 1% Triton-X 100, 500 uM DTT). Lysates were cleared by spinning samples at 17,000 xg for 10 min. Supernatants were transferred to a new 1.5 mL tube. Subunits were split by adding EDTA to 60 mM to samples (mixed in by vortexing) and incubating samples on ice for at least 30 min. Samples were then loaded onto a 10-35% gradient (20 mM Tris-Cl pH 7.5, 140 mM KCl, 5 mM EDTA, 0.5 mM DTT, prepared as described above) and spun at 35,000 rpm for 3.5 h. Traces were obtained by measuring UV absorbance (260 nm) across the gradient using a Biocomp Piston Gradient Fractionator. Traces shown were obtained after normalizing lysates based on cell count. The area under the curve for the 40S and 60S peaks was measured using FIJI.

For sucrose gradients where fractions were used for western blotting, 10%, 20%, 30%, 40% and 50% sucrose buffers were prepared containing 1x gradient buffer. 40 µLs of each buffer were layered onto a 0.2 mL, open-top thickwall polycarbonate tube (Beckmann) in reverse order (50% to 10%). At least 2 µg of RNA were loaded into each gradient in a total volume of 20 µL. Samples were spun for 22 min at 4°C, 50k rpm and were then hand fractionated by taking ten 20µL fractions for each sample.

### Western blots

Samples were prepared by adding 6x Laemmli buffer and boiling for 5-10 min. BCA assays were used to normalize the amount of protein loaded onto gels for whole cell lysates. Samples were loaded onto a 4–12% Criterion™ XT Bis-Tris Protein Gel (Bio-Rad) and run at 150 V for 60 min. Proteins were transferred onto a nitrocellulose membrane using the Trans-Blot Turbo Transfer System (Bio-Rad) high molecular weight setting (10 min). Samples were blocked in 5% milk diluted in 1x TBST for at least 25 min. Primary antibodies (listed below) were added to membranes in 5% milk and incubated overnight at 4°C. Membranes were washed three times for 10 min using 5 mL of 1x TBST and were then incubated in secondary antibodies diluted in 5% milk for at least 1 hour. After a second round of three 10 min washes in 1x TBST, membranes were imaged using a mix of 10% SuperSignal West Femto Maximum Sensitivity Substrate and 90% SuperSignal West Pico PLUS chemiluminescent substrate. The resulting signals were visualized using a ChemiDoc Imaging System (Bio-rad).

| Target | Vendor | Catalogue # | Dilution |
| --- | --- | --- | --- |
| 4EBP-1 | CST | 9644S | 1:1000 |
| alpha-tubulin | CST | 2144 | 1:1000 |
| EDF1 | Abcam | ab174651 | 1:1000 |
| eL33 | ThermoFisher | PA5-103544 | 1:1000 |
| eS19 | Abcam | ab181365 | 1:1000 |
| eS24 | Abcam | ab196652 | 1:1000 |
| eS26 | Proteintech | 14909-1-AP | 1:1000 |
| GAPDH | CST | 2118 | 1:2000 |
| Puromycin | Sigma | MABE343 | 1:1000 |
| RACK1 | CST | 5432 | 1:1000 |
| RIOK3 | Proteintech | 13593-1-AP | 1:1000 |
| RNF10 | Proteintech | 16936-1-AP | 1:1000 |
| SERBP1 | Fortis | A303-938A | 1:2000 |
| uL18 | CST | 51345S | 1:1000 |
| uS3 | Abcam | ab140688 | 1:1000 |
| uS5 | Bethyl | A303-794A | 1:1000 |
| Vinculin | Santa Cruz | sc-73614 | 1:10,000 |

### Mass spectrometry

After fractionation, proteins were isolated from each fraction using cytiva SP3 beads and trypsinized according to the manufacturer’s instructions. Around 50 ng of peptides obtained for each fraction were loaded on the Evotip™ using the standard producer protocol. We utilized EVOSEP coupled with a TimsTOF ULTRA2 (Bruker) mass spectrometer equipped with a Captive Spray II source (Bruker). The separation was carried out on a Performance Column OE measuring 8 cm x 150 µm ID, with a particle size of 1.5 µm, maintained at 40°C. We employed the standard 60SPD EVOSEP method, applying a 21-minute gradient for a total sample-to-sample time of 24 minutes using Solvent A: 0.1% (v/v) formic acid (FA) and Solvent B: acetonitrile (ACN)/0.1% (v/v) FA (Thermo Fisher Scientific™ Optima LCMS grade). The dia-PASEF acquisition scheme was optimized for a cycle time estimate of 0.95 s. The window scheme was designed to cover most of the charge 2 precursor ions in the range m/z 392−1113 and 1/K0 0.67−1.34, using 24 × 25 Th windows, with accumulation and ramp times of 100 ms. The mass spectrometer was operated in the “high sensitivity detection” (“low sample amount”) mode.

### Data analyses

Raw dia-PASEF data were processed with DIA-NN 2.2.0 using a library-free workflow against the human UniProt FASTA database. Identifications were controlled at 1% precursor and protein FDR. DIA-NN processing included match-between-runs. Monosome and polysome fractions were analyzed together, and biological replicates were compared between WT and SERBP1 KO conditions. Analysis of the relative quantities obtained from these searches were performed using a custom R script. Proteins with relative quantities below 500 were excluded from the analyses.

Lists of putative RIH and RIM-containing proteins were generated using SlimSearch4 ^43^ and compared to all proteins included in the mass spectrometry analyses or only to proteins enriched in monosomes or polysomes upon SERBP1 depletion. The RIH search was limited to SΩAX.

Conservation analyses for RNF10 were performed using consurf with default parameters,^44^ while PID calculations were obtained using jalview.^45^

## Author contributions

N.R.-M. and R.G. conceptualized the project and wrote the manuscript. All authors contributed to the experimental design and edited the manuscript. N.R.-M. and E.H.S. performed cell-based experiments and contributed to data analysis. N.R.-M. prepared samples for mass spectrometry and L.S. processed the samples. L.S. and N.R.-M. analyzed the mass spectrometry data.

## Acknowledgements

We are grateful to members of the Green lab for their feedback on experimental design, particularly to Stefan Cehan for technical support and to Joshua Black, Marco Catipovic, Frances Diehl and Eugene Park for carefully reading this manuscript and providing suggestions. This work is supported by a grant from the European Research Council (ERC; to R.G. and J.R.). N.A.R.-M. is supported by an HHMI Hanna Gray Fellowship and a BWF PDEP award.

## Supplemental figure legends

**Figure S1:** A) Western blots for cells transfected with siNT or siSERBP1 and incubated with puromycin using an antibody against puromycin. Ponceau stains are shown as loading control. B) Quantification of lane intensities for puromycin western blots normalized to ponceau stain intensities. C) Western blots for WT and SERBP1 KO cells treated with puromycin using an antibody against puromycin. D) Absorbance traces from sucrose gradient sedimentation of WT or SERBP1 KO cells treated with puromycin for 600 seconds. E) Quantification of harringtonine treatment time course in WT or SERBP1 KO cells. F) Western blot of gradient fractions from untreated or anisomycin (ANS) treated WT cells or from untreated SERBP1 KO cells using antibodies against the ribosome collision marker EDF1 or the ribosomal protein uS3.

**Figure S2:** A) Volcano plot for relative quantities obtained from mass spectrometry showing the log_2_(SERBP1 KO/WT) on the x-axis and the log_10_(p-value) on the y-axis for whole cell lysates from cells treated with DMSO, ANS or TOR. B-C) Scatter plots comparing the log_2_(fold change) of KO/WT for DMSO, ANS and TOR treated monosomes (B) or polysomes (C). Proteins enriched in SERBP1 depleted cells only after ANS treatment are shown in red, those enriched only after TOR treatment are shown in blue and those enriched only in DMSO treated cells are shown in black.

**Figure S3:** A) Absorbance traces from sucrose gradient sedimentation of torin-treated WT or SERBP1 KO cells. A zoomed in view of the 40S and 60S peaks is highlighted. B) Western blots for WT cells transfected with NT siRNAs or WT, RNF10 KO or RIOK3 KO cells transfected with siSERBP1 using antibodies against RNF10, SERBP1, RIOK3, 4E-BP1 (control for mTOR inhibition) or the loading controls vinculin or GAPDH. C) Absorbance traces from sucrose gradient sedimentation of torin-treated SERBP1 KO cells transfected with NT siRNAs or with siRNAs against ASCC3. A closer view of the 40S and 60S peaks is shown.

**Figure S4:** A) Western blots for WT cells transfected with an eGFP control and SERBP1 KO cells transfected with an eGFP control or with different SERBP1 constructs using antibodies against vinculin or SERBP1. B) Western blots of sucrose gradient fractions from SERBP1 KO cells transfected with full length or the ΔmRNA channel binding mutant using antibodies against SERBP1 or the ribosomal protein eS24. C) Absorbance traces from sucrose gradient sedimentation of torin-treated SERBP1 KO cells transfected with plasmids expressing eGFP as a control, full length SERBP1 or SERBP1 mutants where the charge in the region binding the mRNA channel of the 40S subunit have either been neutralized or reversed.

**Figure S5**: A) Predicted aligned error (PAE) plot for RNF10-RACK1 interaction obtained from AlphaFold3. B) Overlay of the structure of SERBP1 and ZAK (PDBs: 9RPV and 9RSX) and the alphafold predictions for RNF10 and USP10 on the ribosomal protein RACK1. C) Western blots for WT cells transfected with an eGFP control and SERBP1 KO cells transfected with an eGFP control or with different SERBP1 constructs using antibodies against vinculin or SERBP1. D) Western blots of sucrose gradient fractions from SERBP1 KO cells transfected with the ΔRACK1 binding mutant using antibodies against SERBP1 or the ribosomal protein eS24. E) Western blots for cells transfected with NT siRNAs or siRNAs against RACK1 for different times using antibodies against RNF10, RACK1 or the loading control alpha-tubulin. F) Western blots from torin-treated WT or RACK1 KO cells transfected with an eGFP control plasmid or with plasmid expressing RNF10 and RIOK3 using antibodies against GAPDH (loading control), RACK1, RNF10 or RIOK3. G) Western blots for WT cells transfected with an eGFP control and for RNF10 KO cells transfected with an eGFP control or different RNF10 mutants or truncations. H) Table comparing the sequences for the FPx(L/F) and S(Ω)Ax(Φ) motifs known or predicted to bind on RACK1 for each of these proteins. Percentage identity (PID) calculations from jalview are displayed in varying shades of purple showing similarities for each residue within the motif across different species for each protein. I) Consurf alignment showing conservation scores for RIM and RIH of RNF10 across ten different species.

**Table S1:** Relative quantities obtained from mass spectrometry experiments performed on untreated, ANS- and TOR-treated wild type and SERBP1 KO HEK-293T cells. Includes the log_2_(KO/WT) values and the p-values calculated for proteins that were differentially enriched upon KO in each condition.

