## Supplementary figures and images for "SERBP1 is a master regulator of ribosome interactions"

### Supplemental Figures

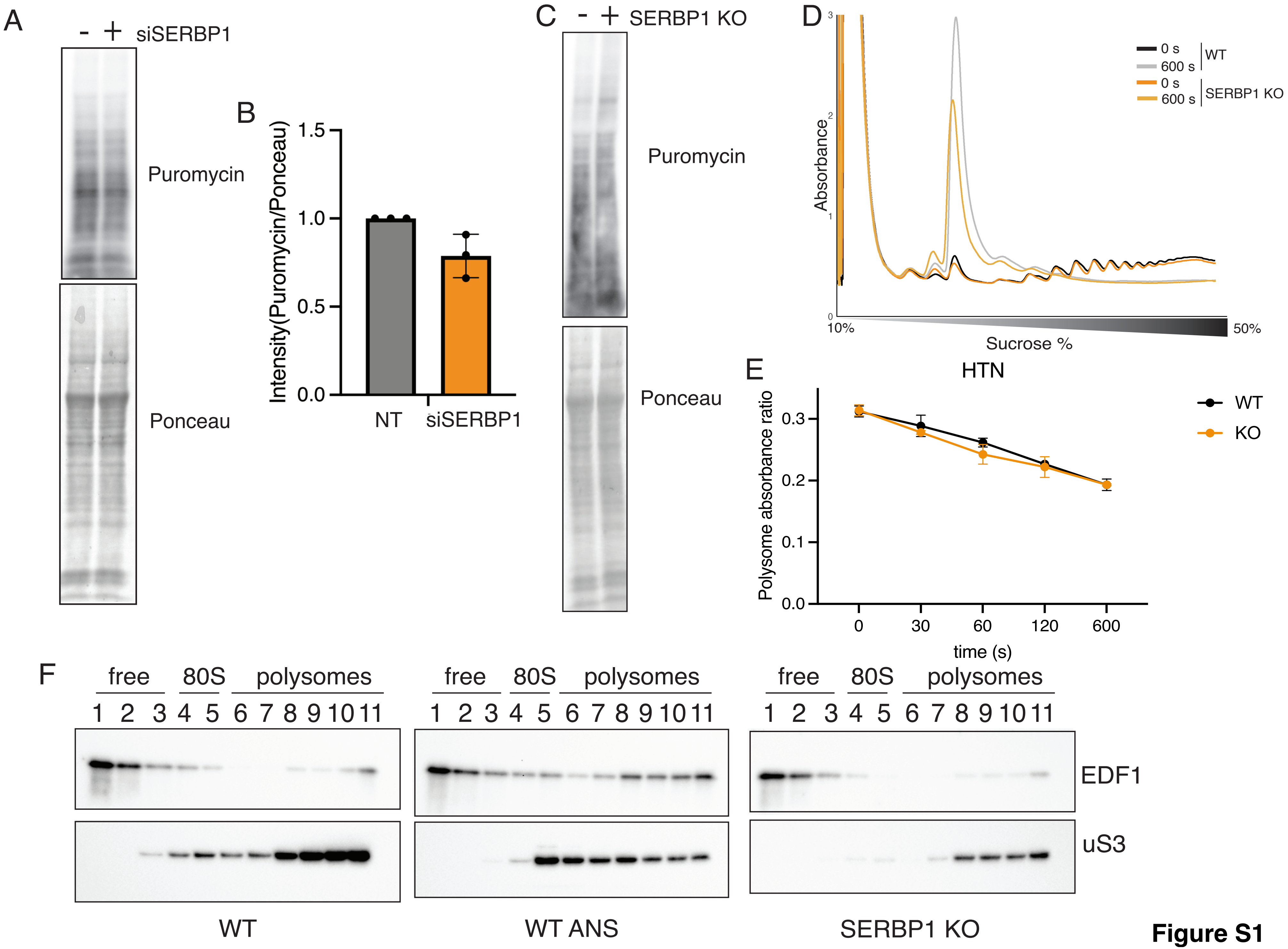

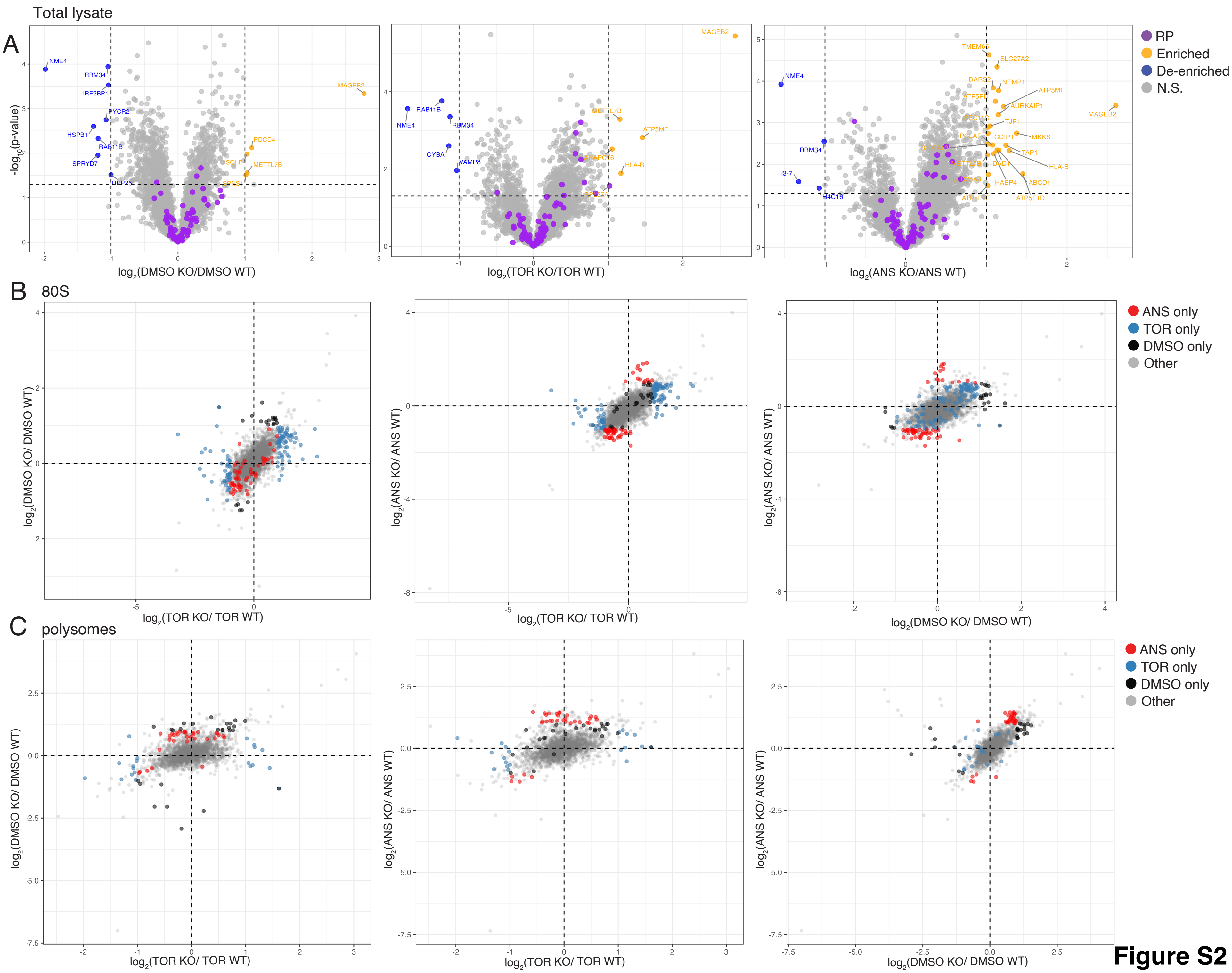

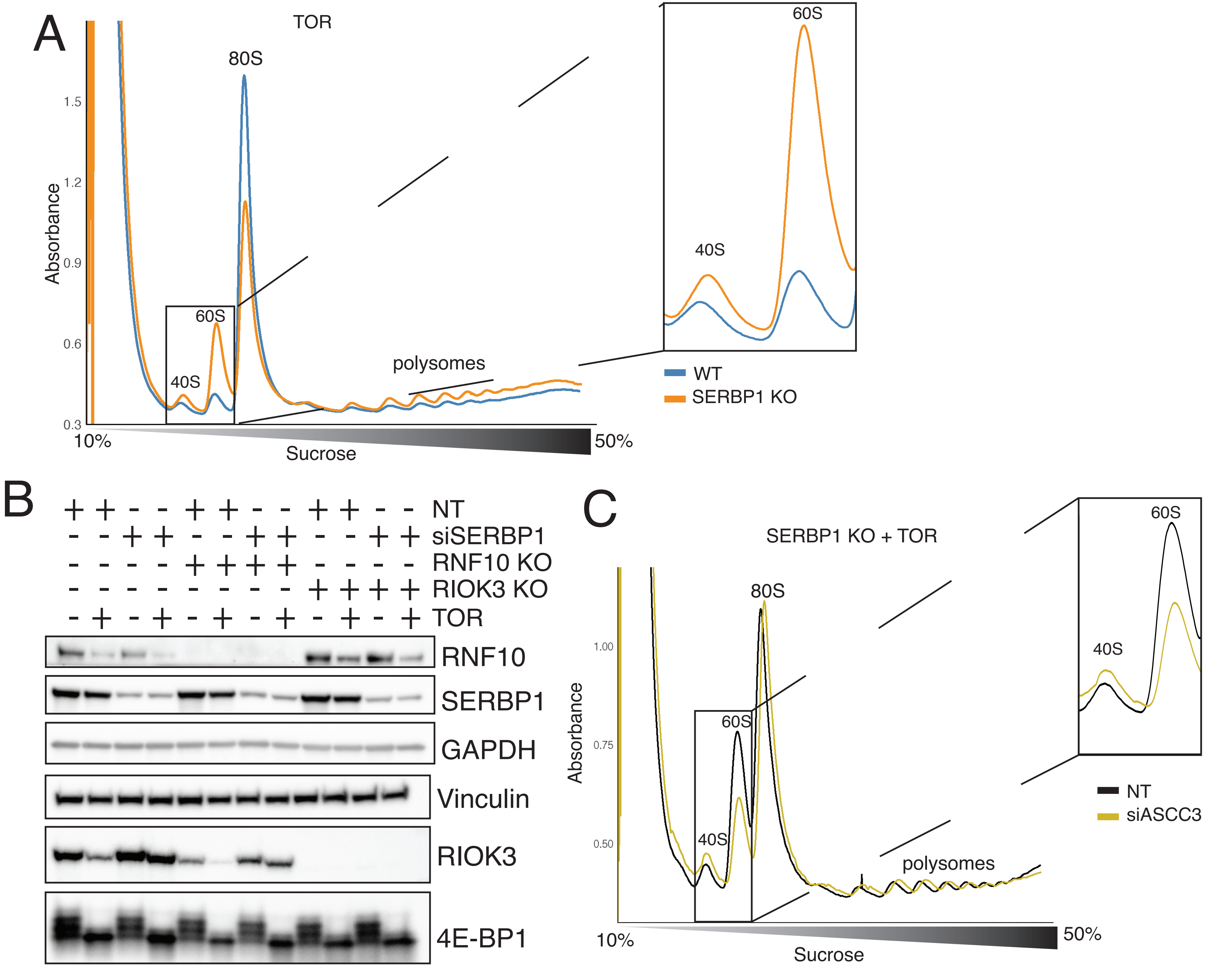

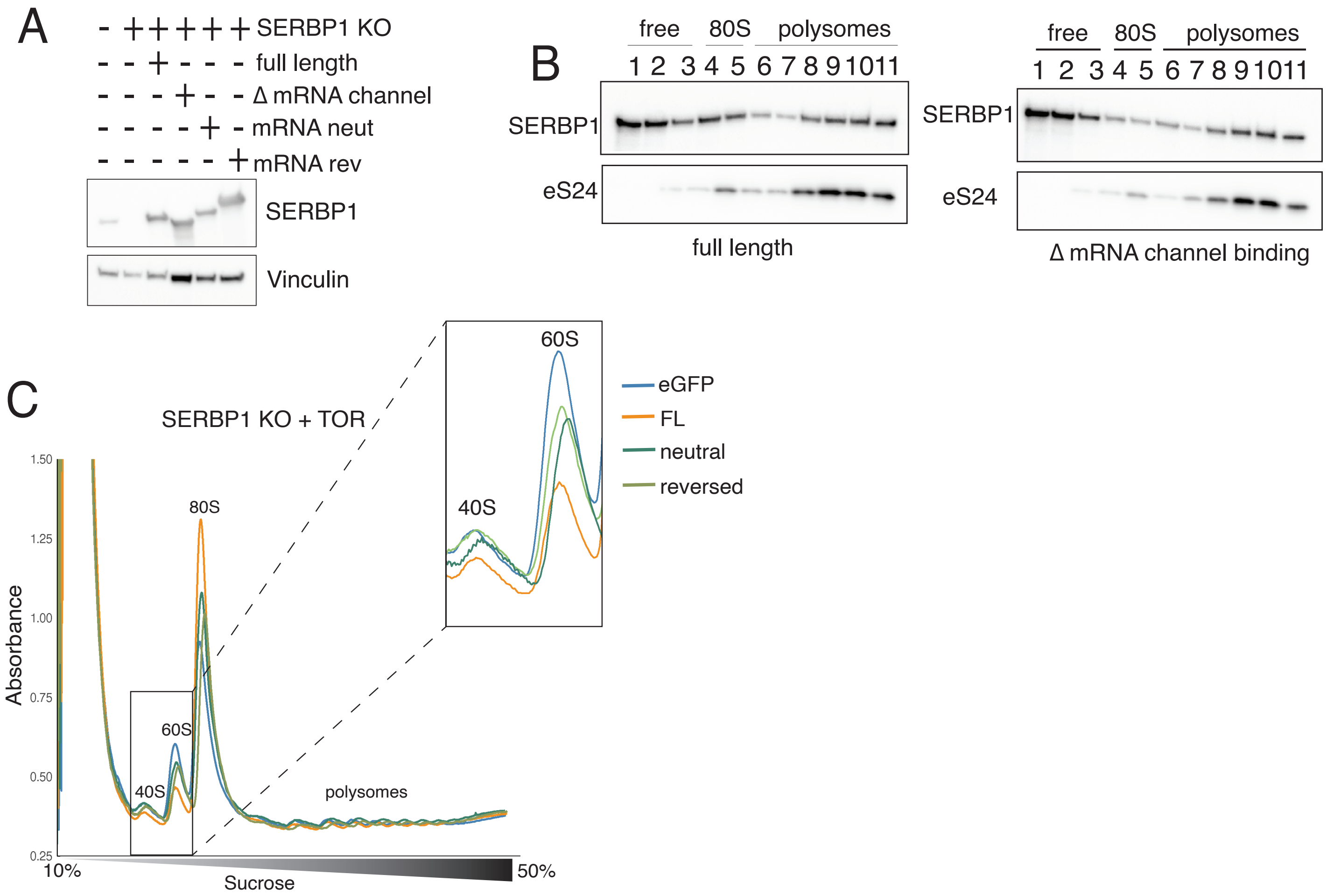

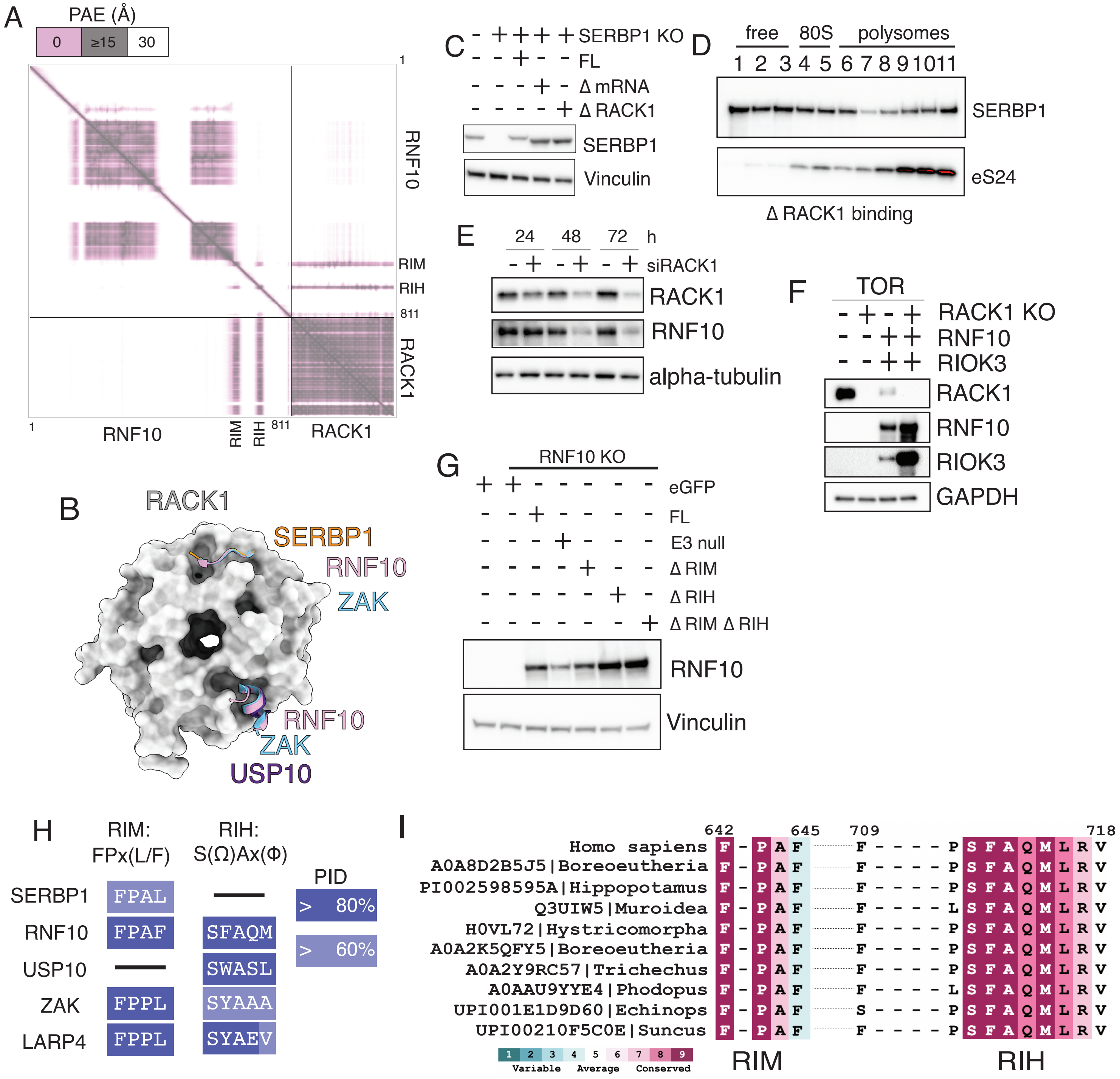
